# Rhythmic information sampling in the brain during visual recognition

**DOI:** 10.1101/2022.06.30.498324

**Authors:** Laurent Caplette, Karim Jerbi, Frédéric Gosselin

## Abstract

When we fixate an object, visual information is continuously received on the retina. Several studies observed behavioral oscillations in perceptual sensitivity across such stimulus time, and these fluctuations have been linked to brain oscillations. However, whether specific brain areas show oscillations across stimulus presentation time (i.e., different time points of the stimulus being more or less processed, in a rhythmic fashion) has not been investigated. Here, we revealed random areas of face images at random moments across time and recorded the brain activity of human participants (both male and female) using magnetoencephalography (MEG) while they performed two recognition tasks. This allowed us to quantify how each snapshot of visual information coming from the stimulus is processed across time and across the brain. Oscillations across stimulus time (rhythmic sampling) were mostly visible in early visual areas, at theta, alpha and low beta frequencies. We also found that they contributed to brain activity more than previously investigated rhythmic processing (oscillations in the processing of a single snapshot of visual information). Non-rhythmic sampling was also visible at later latencies across the visual cortex, either in the form of a transient processing of early stimulus time points or of a sustained processing of the whole stimulus. Our results suggest that successive cycles of ongoing brain oscillations process stimulus information incoming at successive moments. Together, these results advance our understanding of the oscillatory neural dynamics associated with visual processing and show the importance of considering the temporal dimension of stimuli when studying visual recognition.

**Significance Statement:** Several behavioral studies have observed oscillations in perceptual sensitivity over the duration of stimulus presentation, and these fluctuations have been linked to brain oscillations. However, oscillations across stimulus time in the brain have not been studied. Here, we developed a MEG paradigm to quantify how visual information received at each moment during fixation is processed through time and across the brain. We showed that different snapshots of a stimulus are distinctly processed in many brain areas and that these fluctuations are oscillatory in early visual areas. Oscillations across stimulus time were more prevalent than previously studied oscillations across processing time. These results increase our understanding of how neural oscillations interact with the visual processing of temporal stimuli.

## Introduction

Despite seemingly occurring in an instant, visual recognition is a temporal phenomenon. Consider, for a start, a briefly flashed visual stimulus. Its processing by the brain takes a few hundreds of milliseconds. In the real world, however, objects are rarely flashed: they are typically fixated for a few hundreds of milliseconds as well. During this time, their visual information is continuously received on the retina. This second temporal aspect of perception is not acknowledged as often as the first one but is arguably just as important (VanRullen, 2011; King & Wyart, 2021). Depending on the moment at which it is received, information may be processed differently in a given brain area or even not processed at all (Caplette et al., 2020). These variations in processing across stimulus presentation time may be oscillatory, whereby information received at specific moments reoccurring periodically is processed more efficiently (i.e., a rhythmic sampling; Figure 1).

**Figure 1.**
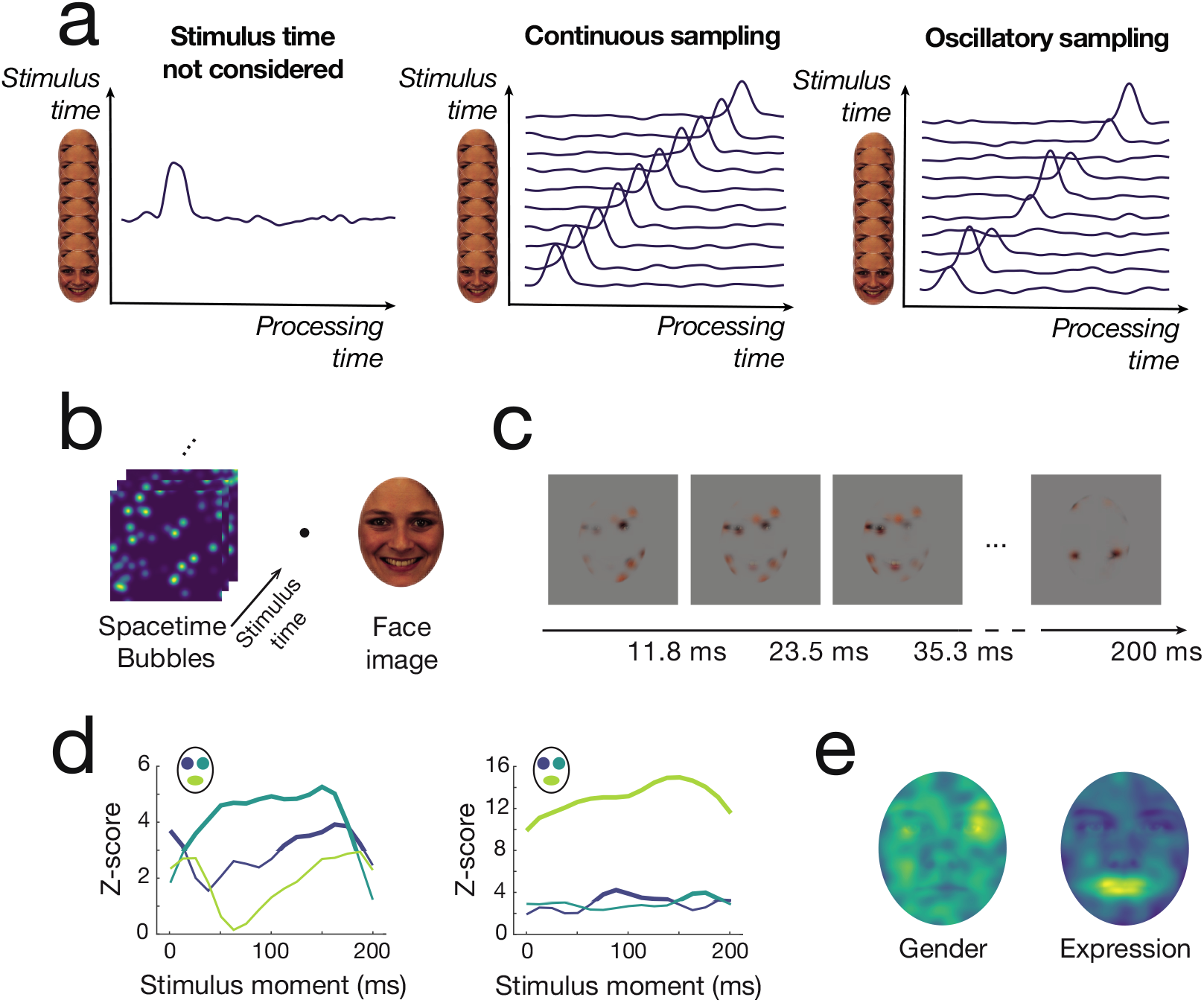
Framework, methods and behavioral results. **a)** Processing of temporally extended visual stimuli in a given brain region. (left) When stimulus duration is not considered, a stimulus is effectively conceptualized as an instantaneous event (only one point on the *y* axis). This visual information is then processed across time (the amplitude of the curve illustrating neural activity). (middle, right) In reality, most stimuli are temporally extended (multiple points on the *y* axis). This entails that information received at each moment during stimulus fixation is processed through time. (middle) Information received at different moments is processed in the same way. (right) Information received at different moments is processed to a different degree (see the varying peak amplitude), in a periodic fashion. **b)** Experimental stimuli were created by dot-multiplying a 3D tensor sparsely populated with 3D Gaussian apertures (Spacetime Bubbles) by a masked face image. **c)** Example stimulus. Random areas of a random face image were smoothly revealed at random moments across 200 ms. See Videos S1-S2. **d)** Time-resolved behavioral group results. Each line depicts how much each face feature presented on each stimulus moment correlated with accurate responses. Bold segments indicate significant correlations (*Z* > 3.12; *p* < .05, one-tailed, FWER-corrected). (left) Gender discrimination task. (right) Expression discrimination task. **e)** Pattern of behavioral group results for the whole face averaged across time. (left) Gender discrimination task. (right) Expression discrimination task.

Previous studies have reported behavioral oscillations in perceptual and attentional sampling across stimulus presentation time (see VanRullen, 2016, for a review). For instance, the visual threshold for detecting two successive flashes oscillates (around 33 Hz) as a function of the time interval between them (Latour, 1967), and the information used to recognize a face seems to be preferentially sampled at a frequency between 10 and 15 Hz (Blais et al., 2013). In addition, when multiple stimuli are monitored simultaneously, attention seems to fluctuate between them at a 7–8 Hz frequency (Dugué et al., 2016; Helfrich et al., 2018; Holcombe & Chen, 2013; Landau & Fries, 2012; Landau et al., 2015; Fiebelkorn et al., 2013; Fiebelkorn et al., 2018; Re et al., 2019; Senoussi et al., 2019; VanRullen et al., 2007; but see van der Werf et al., 2021); this rhythmic sampling seems to occur even when only one object is monitored (Holcombe & Chen, 2013; VanRullen et al., 2007).

These fluctuations have been linked to underlying brain oscillations. Among other findings, detection accuracy has been found to correlate with the phase or power of ongoing or prestimulus theta and alpha oscillations (Busch et al., 2009; Busch & VanRullen, 2010; Hanslmayr et al., 2013; Helfrich et al., 2018; Fiebelkorn et al., 2018) and neural correlates of perception have been found to depend on prestimulus alpha or theta phase (Busch & VanRullen, 2010; Gruber et al., 2014; Hanslmayr et al., 2013; Jansen & Brandt, 1991). However, the neural mechanisms of perceptual rhythmic sampling remain uncertain, with some recent studies having failed to replicate previous findings (Benwell et al., 2017; Buergers & Noppeney, 2022; Chaumon & Busch, 2014; Fekete et al., 2018; Keitel et al., 2022; Ruzzoli et al., 2019; van Diepen et al., 2015). Moreover, sampling (whether oscillatory or not) in the brain has not been directly investigated. That is, it remains unknown whether specific brain areas preferentially process information incoming at specific moments from a stimulus, and whether it can be observed with noninvasive imaging modalities such as MEG. The specific latencies and sampling frequencies at play are also unknown. Finally, if rhythmic sampling is observed, it remains to be seen whether it occurs concurrently with rhythmic processing. Indeed, regardless of whether different stimulus moments are processed differently, visual information could be processed in a rhythmic fashion (e.g., Rousselet et al., 2007; Schyns et al., 2011).

We aimed to investigate these phenomena in the present study. Specifically, we revealed random visual features at random moments across stimulus time, and we correlated neural activity in each brain region and at each time point with the presentation of these features. This allowed us to render and quantify how each visual feature from each stimulus moment is processed across time and across the brain, and thus to investigate both processing and sampling in the brain. We aimed to determine whether rhythmic sampling exists in the brain and, if it does, characterize it in detail and assess its prevalence relative to rhythmic processing.

## Materials and methods

### Participants

Five neurotypical adults (mean age = 27.60; SD = 3.21; 3 female) were recruited on the campus of the University of Montreal; each came to the laboratory for a total of five experimental sessions (5,500 trials per subject were necessary to sample with sufficient signal-to-noise ratio the large search space – all image pixels and stimulus moments – explored in this study). Participants did not suffer from any psychiatric or psychological disorder, had no known history of head concussions, and had normal or corrected-to-normal vision, including color vision. The experimental protocol was approved by the ethics board of the Faculty of Arts and Sciences of the University of Montreal and the study was carried in accordance with the approved guidelines. Written informed consent was obtained from all the participants after the procedure had been fully explained, and a monetary compensation was provided upon completion of each experimental session.

### Materials

The experimental program ran on a Dell Precision computer with Windows XP (Microsoft Inc.) in the Matlab (Mathworks Inc.) environment, using custom scripts and functions from the Psychophysics Toolbox (Brainard, 1997; Pelli, 1997; Kleiner et al., 2007). Stimuli were projected on a screen using a Sanyo PLC-XP41L projector, calibrated to allow a linear manipulation of luminance, with a resolution of 1152 x 864 pixels and an 85 Hz refresh rate. A viewing distance of 55 cm was maintained throughout the experiment. MEG activity was recorded using a 275-sensor CTF scanner with a sampling rate of 1200 Hz. Fiducials at the nasion and left and right temples were used to track head movements in the scanner. Vertical electro-oculogram was bipolarly registered above and below the eyes, and horizontal electro-oculogram was bipolarly registered at the outer canthi of both eyes, to detect blinks and eye movements. An electrocardiogram was used to detect heartbeats. Prior to each session, the surface of the scalp and the positions of the fiducials were digitized using a Polhemus device.

### Stimuli

Two hundred and sixty-four color images of faces were selected from the image database *Karolinska Directed Emotional Faces* (Goeleven et al., 2008); only frontal-view faces were chosen. These were composed of 66 different identities (33 women and 33 men) each performing a happy and a neutral expression; two different pictures of each facial expression were used. Faces were aligned on twenty hand-annotated landmarks averaged to six mean coordinates for left and right eyes, left and right eyebrows, nose, and mouth, using a Procrustes transformation. Face images were then cropped and masked by a centered lightly smoothed elliptical mask to conceal the background, hair and shoulders; final masked faces had a horizontal diameter of 164 pixels, or 6 degrees of visual angle. The mean luminance and contrast of all masked faces were equalized, separately for each color channel, using the SHINE toolbox (Willenbockel et al., 2010).

On each trial (excluding some trials in which an unaltered face image was shown), random areas of a randomly selected exemplar face were revealed at random moments across a total duration of 200 ms (Caplette et al., 2020; Vinette et al., 2004). In other words, different random parts of a face were revealed at different moments during the stimulus. Importantly, the revealed areas were gradually appearing and disappearing (Figure 1c; Videos 1-2). A duration of 200 ms was chosen so that no saccade could occur during stimulus presentation on most trials and because it represents the duration of a typical eye fixation. There were 17 frames (hereafter referred to as *stimulus moments*) each lasting 11.76 ms. To create such a stimulus, we first generated a random 480 (pixels) × 480 (pixels) × 25 (stimulus moments, including padding that was not used for the final stimulus) sparse tensor composed of 0s and a few 1s. The probability of each element being 1 was adjusted on a trial-by-trial basis using a gradient descent algorithm to maintain accuracy around 75%. Three-dimensional Gaussian apertures (i.e., Bubbles; α_space_ = 11.4 pixels, or 0.42 degree of visual angle; α_time_ = 22.3 ms) (Gosselin & Schyns, 2001; Vinette et al., 2004) were then centered on each 1. On each trial, there was a minimum of one bubble in the (non-padding portion of the) stimulus (the average number of bubbles was 94.74). Superfluous padding was then removed so that the tensor was 320 (pixels) × 320 (pixels) × 17 (stimulus moments), and thresholding was applied so that no value exceeded one. All seventeen 320 × 320 matrices composing a tensor were finally multiplied elementwise with the same elliptical face mask described above. We called this 3D array *sampling tensor* and the value of each element determined the visibility of a given pixel in a given stimulus moment for this trial. Specifically, the face image (repeated across the temporal dimension) and the sampling tensor were multiplied elementwise together, the complement of the sampling tensor was multiplied elementwise with a mid-grey plane and both results were summed together, so that non-sampled parts of the face were replaced by a median grey. Note also that on each trial, the underlying face image had a 50% probability of being flipped along the vertical midline, to compensate for possible informational differences between the left and right sides of the face images used.

### Experimental design and statistical analyses

#### Experimental design

Each participant came to the MEG Laboratory of the Department of Psychology, University of Montreal, five times. They filled a personal information questionnaire (education, age, sex, hours of sleep, alertness, concussion history, mental illness history, etc.) on the first session. Participants completed between 1,000 and 1,350 trials on each session, in blocks of 250 or 275 trials separated by short breaks. On each block, participants performed either a gender discrimination task (“man or woman?”) or an expression discrimination task (“happy or neutral?”). Tasks alternated on each successive block and the first task performed was counterbalanced across subjects. Overall, each participant completed 2,850 trials of each task (except 2,875 in one case): 2,750 Bubbles trials (see above) and 100 trials (125 in one case) in which whole non-sampled faces were shown. Whole-face trials per task were randomly intermixed with Bubbles trials within four blocks (five in one case) of each task (participants had to perform the same task and faces were also shown for 200 ms). Whole-face trials were not analyzed in this manuscript and, unless explicitly stated, all further statements refer only to Bubbles trials. Participants were instructed not to blink during the trials themselves. After every 5 trials, the screen automatically showed text indicating to the participants that they could take a few seconds to blink and rest their eyes before pressing a key to continue the experiment.

On each trial, a central black fixation cross was shown to the participants for a constant 1500 ms, after which the video stimulus appeared during 200 ms, superposed on the fixation cross, again followed by the fixation cross until the participant responded; a mid-grey background was always present. A fixed interval (1500 ms) between the response and the next stimulus was used so that participants could predict the onset of the trials. Participants had to respond as accurately and rapidly as possible with two keys on the keyboard (key-response associations were counterbalanced across participants).

#### Behavioral data analysis

We aimed to analyze how each face feature shown at each moment during the stimulus led to more or less accurate responses. To do so, we first excluded trials with a response time +/– 3 standard deviations away from the mean response time, below 100 ms or above 2,000 ms from further analyses (1.37% of trials). Accuracies were then transformed in z-scores across trials for each session. Sampling tensors were downsampled across the spatial dimensions to 64 × 64 pixels and transformed in z-scores across trials. Sampling tensors and accuracy vectors of all sessions were concatenated for each subject and task. An outer product was then performed between the vectorized sampling tensors and the accuracies for all trials (given the random sampling, this is equivalent to a multiple linear regression). This results in a *classification volume* indicating how each pixel at each moment correlates to accurate responses. The analysis was then repeated while randomly permuting the accuracies 1,000 times to establish a null distribution of classification images, and results were averaged across subjects. To observe the pixel-resolution time-averaged pattern of correlations, all 17 stimulus moments were averaged together, and the observed results were z-scored using the distribution of null results. To observe the time-resolved patterns of correlations for the main face features, pixel ROIs were created using lightly smoothed circles (for the eyes) and ellipse (for the mouth). All three ROIs contained a similar number of pixels (within a 1% margin). A scalar product was then performed between the masks of the ROIs and the classification images. Finally, the observed and null results were transformed in z-scores using the distribution of null results, and the maximum statistic method was applied to establish a statistical threshold while correcting for multiple comparisons (*p* < .05, one-tailed, FWER-corrected; Holmes et al., 1996).

#### MEG preprocessing and source reconstruction

Participants had all been MRI scanned in various studies over the course of the past five years; these anatomical T1 MRIs were used for source reconstruction in the current study. All MRIs were obtained using a MP-RAGE sequence on a 3T Siemens (Trio or Prisma) scanner with a 1 mm × 1 mm × 1 mm spacing.

The cortical surface was extracted using Freesurfer (http://surfer.nmr.mgh.harvard.edu) and downsampled to 8,000 vertices using Brainstorm. All further preprocessing was conducted using functions from the Brainstorm toolbox (Tadel et al., 2011). Anatomical landmarks corresponding to the locations of the MEG fiducials were identified and the MRI was aligned with the MEG sensors. The digitized surface of the scalp was used to refine the coregistration.

When necessary, MEG runs were subdivided so that the head did not move more than 3.5 mm within each analyzed brain activity segment; short segments with large head movements were also excluded in the process. Data from all resulting segments were band-passed between 1 and 40 Hz (Coquelet et al., 2020; Zhan et al., 2019) using a Kaiser FIR Filter with 60 dB stopband attenuation and resampled to a 250 Hz sampling rate to reduce data dimensionality. Bad channels were visually identified and removed. Signal Space Projection (SSP; Tesche et al., 1995) was used to identify and remove remaining saccade, blink and heartbeat artifacts. When SSP failed to identify artifacts, an ICA and correlations of component time courses with ECG or EOG time courses were used. The data were then segmented into trials from –250 ms to 600 ms from the stimulus onset, and baseline corrected using the average activity between 250 ms and 0 ms before the stimulus onset. Trials with noisy or anomalous segments, or with blinks/saccades during or close to the stimulus, were removed following a visual inspection. Like for the behavioral data analyses, trials with a response time +/– 3 standard deviations away from the mean response time, below 100 ms or above 2,000 ms, were also excluded. In sum, 590 trials were excluded (2.15% of all trials) and 26,910 trials entered subsequent analyses.

For each run, a forward head model was constructed using overlapping spheres (Leahy et al., 1998). A noise covariance matrix was then estimated from (filtered and resampled) empty-room noise recordings from the same day and it was regularized using optimal shrinkage (Ledoit & Wolf, 2004). Estimation of the activity at 8,000 points of the cortical surface was performed for each trial using the minimum norm technique. One dipole, oriented normally to the cortical surface, was estimated at each vertex. Source maps were normalized using dynamical Statistical Parametric Mapping (dSPM; Dale et al., 2000).

#### Creation of stimulus time × MEG time maps

We aimed to uncover how much each face feature shown at each moment during the stimulus was processed at each brain source. MEG activity was first transformed in z-scores across non-excluded correct trials for each session, source and time point. Pixel ROIs for the three main face features were used to create sampling matrices (3 face features × 17 stimulus moments) from the 3D sampling tensors (see Behavioral data analysis). Sampling matrices of all trials were then vectorized. The resulting 51 elements were z-scored across all correct trials; they were then concatenated across all sessions, separately for each task. Source activity for all correct trials was also concatenated in the same way.

For each task, source, and time point, a ridge regression was performed between elements of the sampling matrices (indicating the visibility of each face feature at each stimulus moment) and MEG source activity of all correct trials, using a regularization parameter of 20,000 (Figure 2a). Regression coefficients were then transformed into absolute values, and Gaussian-smoothed across the cortical surface (α_cortex_ = 2.5 mm) and across time (α_time_ = 12 ms). Analyses were repeated 250 times while randomly permuting sampling matrices across trials to establish an empirical null distribution, and the observed regression coefficients were z-scored using the distribution of null regression coefficients. To visualize the sampling and processing patterns, we rearranged these regression coefficients into one stimulus time × MEG time map for each subject, task, face feature and cortical source.

**Figure 2.**
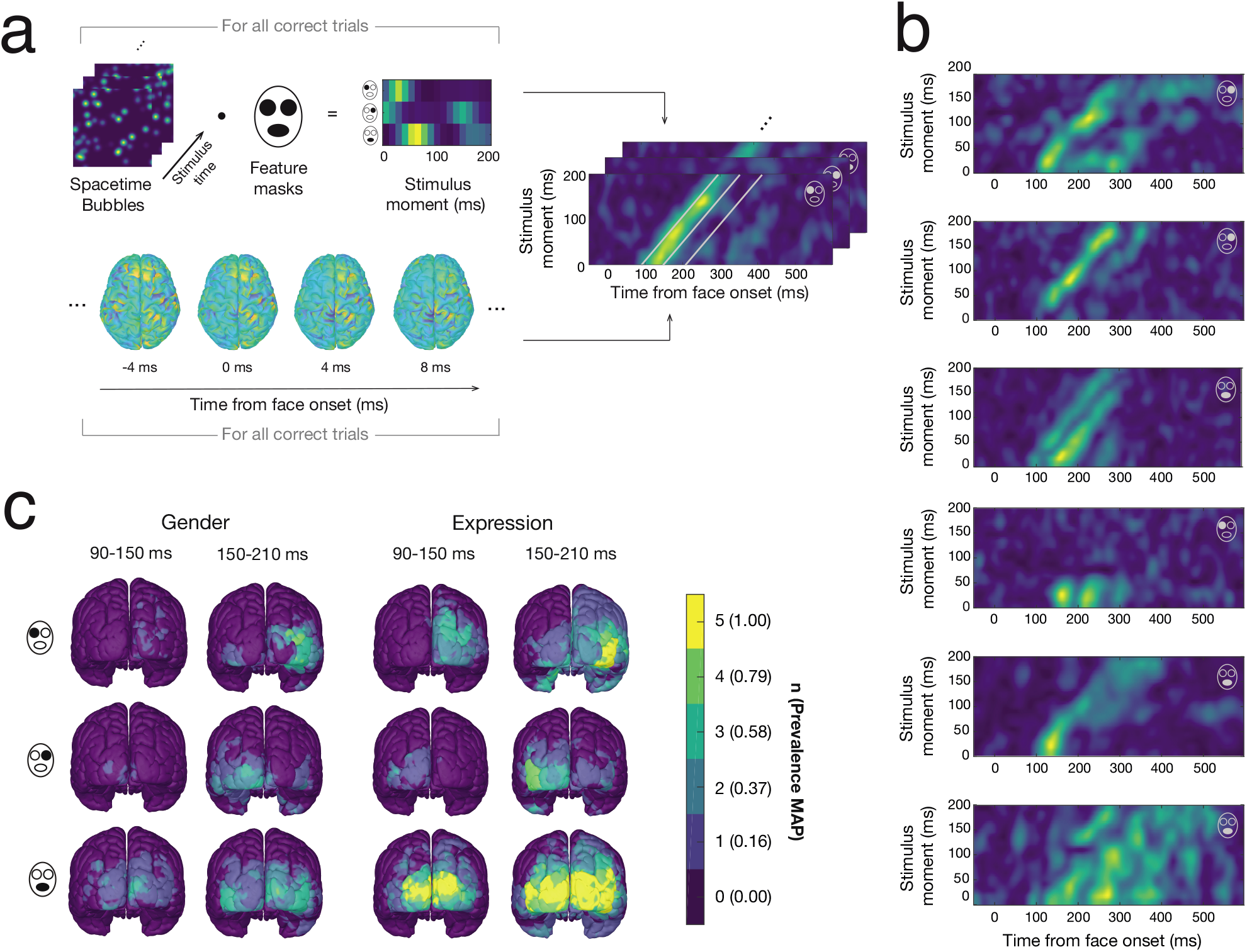
Characterizing sampling and processing across the brain. **a)** Regression analysis. Stimulus dimensionality is first reduced from the full image plane to the 3 main face features (the stimulus time dimension is unaltered). Then, ridge regressions are performed to uncover how each face feature on each stimulus moment (independent variables) explains brain activity at each cortical source and time point (dependent variables) on correct trials: the result is one stimulus time × MEG time map for each participant, cortical source, task, and face feature. Each map indicates how visual information received at each moment on the retina (*y* axis) is processed across time (*x* axis). The time windows of interest for further analyses (90– 150 ms and 150–210 ms from the feature onset) are illustrated by the diagonal grey lines in the top map. **b)** Example maps are depicted (from top to bottom, these maps correspond to: (i) S1, gender task, right eye, left occipital pole; (ii) S3, gender task, right eye, left occipital pole; (iii) S3, gender task, mouth, left middle occipital gyrus; (iv) S4, gender task, left eye, right inferior temporal gyrus; (v) S4, expression task, mouth, right calcarine sulcus; (vi) S5, expression task, mouth, right middle occipital gyrus). Lighter colors indicate greater processing. The face feature associated to a given map is depicted by a grey face icon in the top right corner of the map. **c)** Variance in the processing of each face feature in each task across stimulus moments, in two different time windows. Note that variance is computed in diagonal time windows on the time × time maps (i.e., take the larger delay from later stimulus moments into account; see panel a). The color scale refers to the number of significant participants (within-subject randomization test, *p* < .05, one-tailed, FWER-corrected) and the Bayesian MAP of the estimated population prevalence.

#### Analysis of activity variance across stimulus moments

We aimed to analyze whether the processing of a given face feature at a given latency was different depending on when it was shown during the stimulus. We first averaged, for each stimulus moment, the regression coefficients between 90 and 150 ms, and between 150 ms and 210 ms, from the *feature onset*. These latencies refer to the *x* axis on the time × time maps but they are relative to the feature onsets rather than the face onset, i.e., they take into account the expected processing delay of information shown at later stimulus moments, and so the resulting time windows are diagonal on the time × time maps (see Figure 2a). Note that these specific time windows were chosen because they include the latencies of the two main visual evoked potentials in response to faces, the P1 and the N170, and therefore they are likely to contain most stimulus-related brain activity (Luck, 2012; Mangun, 1995; Rossion & Jacques, 2012). We then computed the variance of these averaged coefficients across the 17 stimulus moments (*y* axis on the time × time maps), for each time window. We repeated these analyses with the permutation maps to obtain a null distribution of variance. The observed and null results were transformed in z-scores using the distribution of null results and they were projected onto the MNI template brain (15,000 vertices). For each subject, a statistical threshold (*p* < .05, one-tailed, FWER-corrected) was determined using the maximum statistic method (Holmes et al., 1996). We then computed a Bayesian estimate of the prevalence of a significant effect in the population for each source, task and feature (default uniform prior and parameters). Given the sample size, the number of participants with a significant effect, and the alpha of the within-subject significance test, this method computes the most probable prevalence of the effect in the population (maximum a posteriori, MAP) and a confidence interval around that value (highest posterior density interval, HPDI) (Ince et al., 2021, 2022). This approach enables a robust population-level statistical inference with a small number of participants. Since a separate statistical test is performed in every participant, each participant acts in effect as a separate replication. Moreover, within-subject effect sizes are not used in the estimation of an overall effect size, preventing its artificial inflation common in small samples (Picciotto, 2020). Lastly, results were projected to a higher-resolution template brain (306,716 voxels) for visualization (this is also the case for all other analyses).

#### NMF and clustering analyses

We then aimed to describe the different types of activity patterns visible across the brains of participants. We first reduced the large number of maps to a few representative exemplars for each subject, task and face feature using nonnegative matrix factorization (NMF). The NMF algorithm requires the *a priori* setting of a *k* parameter representing the number of components to fit. To choose the value of this parameter *k*, (i) we divided the trials in two random halves and repeated the regression analyses described above for each half to obtain two sets of time × time maps; (ii) for each possible parameter *k* between 1 and 12, we computed a R^2^ value measuring how well the components extracted by the NMF algorithm on one half could reconstruct the original data on the other half; (iii) we repeated this analysis for both halves and averaged the results; (iv) we chose the *k* value that maximized the average R^2^. We then applied the NMF on the complete dataset using the chosen *k* value for each subject, task, and feature. The weights thus obtained were finally reapplied to the observed maps and applied to the null maps, and both resulting observed and null NMF maps were transformed in z-scores using the null NMF maps. The resulting NMF maps are the maps depicted on Figure 3. A statistical threshold that corrects for multiple comparisons was established using the maximum statistic method (*p* < .05, one-tailed, FWER-corrected; Holmes et al., 1996).

**Figure 3.**
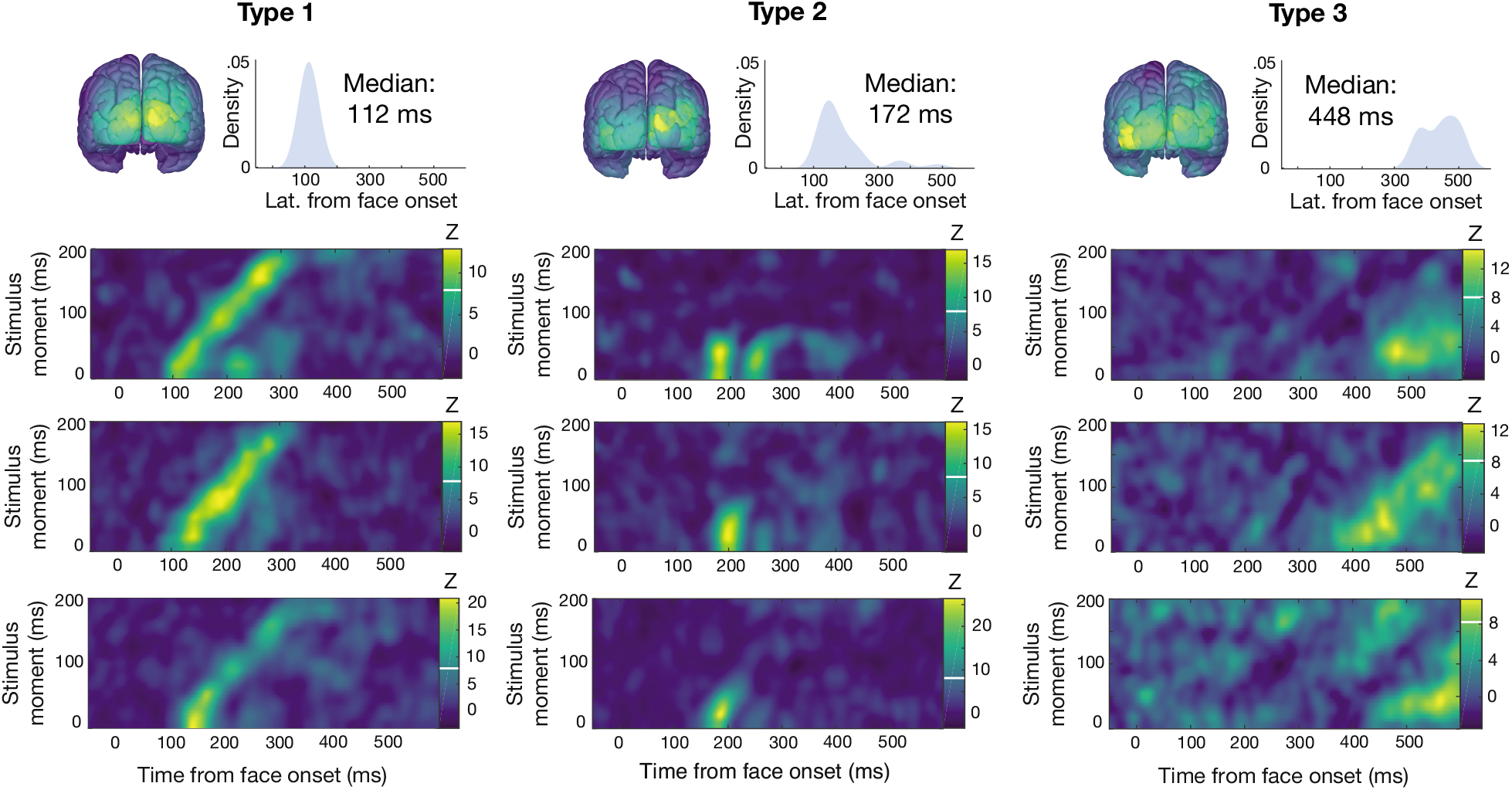
Types of sampling patterns, assessed as clusters in high-dimensional pattern space. For each type, the average brain location, the distribution of onset latencies (with median latency) and example NMF maps are depicted. White horizontal bars indicate the statistical significance threshold (*Z* > 8.12; *p* < .05, one-tailed, FWER-corrected across maps of all subjects) of the activity in the maps. Type 1 maps correspond, from top to bottom, to: S1, gender task, left eye; S3, gender task, mouth; S1, expression task, mouth. Type 2 maps correspond to: S4, expression task, left eye; S1, expression task, mouth; S3, gender task, mouth. Type 3 maps correspond to: S4, expression task, mouth; S5, expression task, mouth; S5, expression task, mouth.

To identify types of NMF maps, we performed an exploratory clustering analysis, using the fast search and find of density peaks algorithm (Rodriguez & Laio, 2014). All maps resulting from the NMF analysis that had at least some significant activity were given to the clustering algorithm after reducing their dimensionality from 2,788 (164 time points × 17 stimulus moments) to 24 (the number of principal components necessary to explain at least 90% of the variance) using PCA. The clustering was computed using the correlation distance between the maps to ensure that the global magnitude of each map was not considered. The points on the decision graph that had a *rho* value larger than 1 and a *delta* value two standard deviations above the mean were selected to form clusters. This resulted in three clusters. The latency distribution of each cluster was computed using the first latency above significance for each map and a Gaussian kernel with a fixed bandwidth of 30 ms.

#### Analysis of oscillatory sampling and model fit

We then investigated oscillations in sampling – that is, oscillations across the stimulus time dimension in the stimulus time × MEG time maps (Figure 4b). Specifically, we used regression coefficients neither smoothed across the cortical surface nor across time, and averaged coefficients within specific time windows (90−150 ms and 150−210 ms from the feature onsets, same windows as in the previous variance analysis; Figure 2a). For each source and face feature, resulting activity across all 25 (stimulus + padding) stimulus moments was linearly detrended, windowed and Fourier transformed. Given the short signals, we used a Tukey window with a taper value of 0.1. Fourier coefficients were then smoothed across the cortical surface (α_cortex_ = 2.5 mm). Frequencies from 6.8 Hz to 30.4 Hz were analyzed; lower and higher frequencies could not be analyzed due to the stimulus length, resolution and temporal smoothing. These analyses were repeated on the null distribution regression coefficients. Both observed and null distribution Fourier coefficients were z-scored with the null distribution Fourier coefficients and projected onto the MNI template brain. Results were averaged across tasks and face features. For each subject, a statistical threshold that corrects for multiple comparisons (*p* < .05, one-tailed, FWER-corrected) was determined using the maximum statistic method (Holmes et al., 1996). We then computed a Bayesian estimate of the population prevalence of a significant effect for each source, time window and frequency (default uniform prior and parameters; Ince et al., 2021). Note that our empirical null distribution has the same aperiodic temporal structure as our observed data (unlike if we were to use a time-shuffling method) and therefore our statistical threshold should not be liberal for any frequency (see Brookshire, 2022). Moreover, since we normalize our results with the null distribution for each frequency separately, any potential artifact or non-uniform spectrum resulting purely from our method (also appearing in the null distribution) is removed from the data.

**Figure 4.**
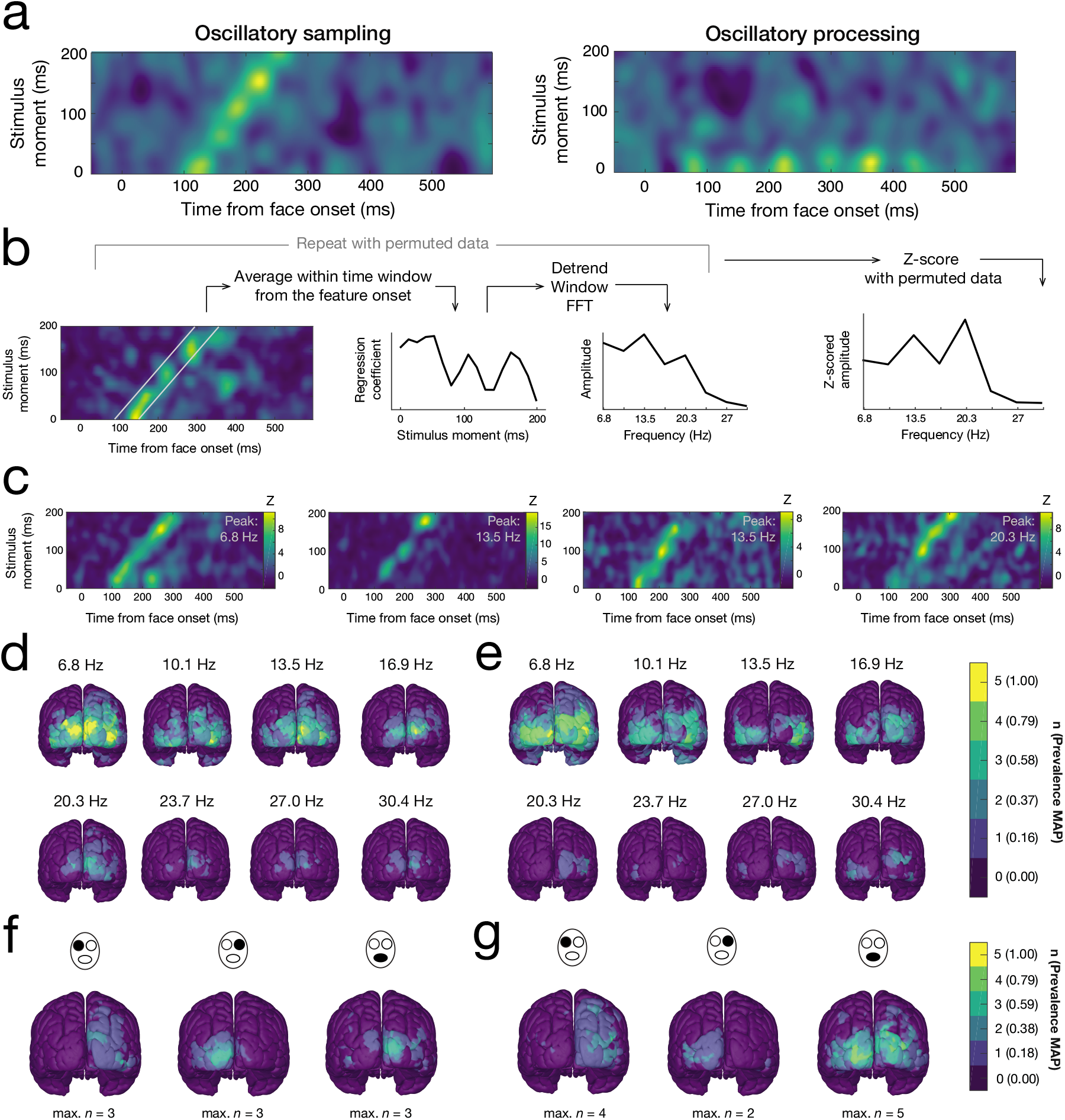
Rhythmic sampling across the brain. **a)** Simulated data illustrating the difference between sampling oscillations (left) and processing oscillations (right). Note that each map depicts only one possible manifestation of the oscillatory phenomenon illustrated. **b)** Sampling oscillations analysis. Activity is first averaged within a diagonal time window to obtain one coefficient per stimulus moment. This series of regression coefficients (including padding, not illustrated) is then detrended, windowed and Fourier-transformed. This analysis is repeated with the null distribution of maps and the observed Fourier coefficients are z-scored with the null Fourier coefficients. **c)** Example maps with significant sampling oscillations. The sampling frequency with the highest amplitude is indicated in the top right corner. **d-e)** Amplitude of sampling oscillations at each frequency, across the brain, averaged across tasks and face features, for the 90–150 ms time window (d) and the 150–210 ms time window (e). The color scale refers to the number of significant participants (within-subject randomization test, *p* < .05, one-tailed, FWER-corrected) and the Bayesian MAP of the estimated population prevalence. **f-g)** Sampling oscillations vs processing oscillations model fit, for each face feature, for the gender (f) and expression (g) tasks. The color scale refers to the number of participants for which the sampling oscillations model fit better (within-subject randomization test, *p* < .05, two-tailed, FWER-corrected) and the Bayesian MAP of the estimated population prevalence. The maximum number of participants across the cortex is indicated below for each feature and task. A better fit of the processing oscillations model never occurred for more than one participant (see Extended Data Figure 4-1).

To test whether sampling oscillations explained activity better than processing oscillations, we fitted two models to each time × time map: one representing oscillations across the horizontal processing time dimension (i.e., processing oscillations) and one representing oscillations across the vertical stimulus time dimension (i.e., sampling oscillations). To equalize the data on which both models were fit and to account for delays associated with later stimulus moments, the models were fit on maps in which activity for later stimulus moments was shifted left proportionally to their expected processing delay (because of this necessary shift, only 400 ms of processing were included for each stimulus moment). Each model was created by summing seven sine waves of frequencies from 6.76 Hz to 27.04 Hz in one dimension and repeating the resulting wave across the other dimension. The magnitudes and phases of each sine wave were free parameters (14 free parameters for each model). Best-fit magnitudes were estimated using FFTs (using the same detrending and windowing procedure as above) across the relevant dimension. For the phases, initial values were first estimated using FFTs across the relevant dimension and the models were then fitted 100 times using random values pi radians around these initial values. For each model, the values that minimized mean squared error were chosen. To compute a measure of model fit, the final mean squared errors were smoothed across the cortical surface (α_cortex_ = 2.5 mm) and the difference between the models was computed. This whole procedure was then repeated with the null time × time maps. For each subject, task, face feature, and cortical source, the observed and null final difference scores were z-scored using the null final difference scores. Results were finally projected onto the MNI template brain, and statistical thresholds that correct for multiple comparisons (*p* < .05, two-tailed, FWER-corrected) were established using the maximum statistic method (Holmes et al., 1996) for each subject. We then computed a Bayesian estimate of the population prevalence of a significant effect for each source, task and face feature (default uniform prior and parameters; Ince et al., 2021).

#### Code accessibility

All custom scripts and functions relevant to this study are available in an Open Science Framework online repository (https://osf.io/pcybk/).

#### Data availability

All raw and preprocessed data pertaining to this study is available in an Open Science Framework online repository (https://osf.io/pcybk/).

## Results

We recorded the brain activity of five neurotypical adults over five days each with MEG. On each trial, we showed face images to the participants, and they had to categorize either the gender (man vs woman) or the facial expression (happy vs neutral) of the face in alternating blocks of trials. Importantly, each face image was only partially visible: we revealed random parts of each face image at random moments across the 200 ms stimulus duration (Figure 1b-c; Videos 1-2; Caplette et al., 2020; Vinette et al., 2004). Accurate categorizations mostly correlated with the presentation of the eyes in the gender discrimination task, while they mostly correlated with the presentation of the mouth in the expression discrimination task, and these correlations were largely maintained throughout stimulus presentation, with some fluctuations (*Z* > 3.12; *p* < .05, one-tailed, FWER-corrected, randomization test; Figure 1d-e). These results replicate previous findings (Dupuis-Roy et al., 2009, 2019; Faghel-Soubeyrand et al., 2019; Gosselin & Schyns, 2001; Schyns et al., 2002).

### Characterizing sampling and processing across the brain

Stimulus time × MEG time maps were obtained for each subject, task, face feature and cortical source (Figure 2a). These maps reveal an as-of-yet unseen dimension of visual processing: they show whether and how information received on the retina at different moments throughout stimulus presentation (i.e., *stimulus moments*) is simultaneously processed through time in a specific brain area. By visualizing both stimulus time and MEG time simultaneously, we can untangle the brain activity due to visual information received at different moments, and more fully characterize the dynamics of visual recognition in the brain (see examples of maps on Figure 2b).

In most of these maps, information seemed to be processed in a highly variable manner across stimulus moments. We quantified this variance in brain activity for all tasks, face features and brain sources, looking at diagonal time windows where most activity occurred: 90–150 ms and 150–210 ms from the feature onset (*x* axis on the time × time maps, diagonally to account for the expected processing delay of each stimulus moment; see Figure 2a). There was significant variance for at least three participants (population prevalence MAP = 0.58, 96% HPDI = [0.18, 0.90]; within-subject randomization test, *p* < .05, one-tailed, FWER-corrected) in all cases except for the eyes in the gender task in the first time window, for multiple areas across the occipital, temporal and parietal lobes, and for both time windows investigated. Most notably, all participants showed significant variance across stimulus moments for the processing of the mouth (both time windows) and the left eye (late time window) in the expression task, indicating that at least 56% (and most likely 100%) of the population would likely show this effect (population prevalence MAP = 1.00, 96% HPDI = [0.56, 1.00]). Significant variance generally peaked in the occipital poles early on, and then extended to the inferior and middle occipital gyri. These findings confirm that the exact moment at which information is received on the retina matters for its processing in the brain. This variance could be caused by many different underlying processes, including oscillatory sampling, temporal integration, and top-down visual routines (Caplette et al., 2020).

We characterized the different kinds of activity patterns that were occurring across the brains of the different subjects. To do so, we first reduced the number of maps for each subject, task and face feature using nonnegative matrix factorization (NMF). We tested the significance of the activity in these NMF maps and confirmed that significant processing (*Z* > 8.12; *p* < .05, one-tailed, FWER-corrected, randomization test) occurred in multiple brain areas, at different latencies and for various stimulus moments, for all subjects (see example maps on Figure 3). We then input all NMF maps with significant activity to a clustering algorithm (Rodriguez & Laio, 2014) to identify reliable types of maps, irrespectively of potential small differences in latencies or amplitudes. We identified three types of patterns. In maps of the first type, a clear diagonal trend was visible in the activations across stimulus moments, meaning that information received at all moments was more or less always processed with a constant delay after its reception. However, information from different stimulus moments was processed differently: information received at some moments elicited a greater activity than information received at other moments, in a somewhat rhythmic fashion (see also next section). This pattern was mostly visible in early visual areas (peaking in the occipital poles) and at early latencies (median of 112 ms) (Figure 3, left). It is consistent with the idea that the earliest brain areas simply process all incoming sensory information in an automatic bottom-up fashion. Another type of pattern was visible in partly overlapping areas (occipital poles) but also extending to further loci (left inferior occipital gyrus and right middle occipital gyrus) and later latencies (median of 172 ms). In maps of this type, only information received in the first 50–75 ms of stimulation was processed, most often in one brief burst, and sometimes in two successive bursts (Figure 3, middle). Preferential processing of these stimulus moments may occur because the information received early is generally prioritized, possibly because the total amount of useful information to come is uncertain (Nienborg & Cumming, 2009). The third and final cluster includes maps that show late (median of 448 ms) and relatively long-lasting processing of most stimulus moments. This pattern can be found in slightly higher-level brain areas on average, peaking in the left inferior occipital gyrus and extending up to the right parietal cortex (Figure 3, right). These maps likely represent a higher-level processing of sensory information that is occurring in later areas related to object recognition and evidence accumulation (Hanks et al., 2015; Woolnough et al., 2020).

### Rhythmic sampling across the brain

We then investigated oscillations in sampling – that is, oscillations across the stimulus time dimension in the stimulus time x MEG time maps (Figure 4a-b). We focused on activity in the same two diagonal time windows as for the variance analysis (90–150 ms and 150–210 ms after feature onsets; see Figure 2b). Looking first at the earlier time window, we uncovered significant oscillations in several participants at most analyzed frequencies. Remarkably, all participants showed significant oscillations at all frequencies from 6.8 Hz to 16.9 Hz (population prevalence MAP = 1.00, 96% HPDI = [0.56, 1.00]; within-subject randomization test, *p* < .05, one-tailed, FWER-corrected). These oscillations were observed mainly in the occipital poles, cunei, inferior occipital gyri and middle occipital gyri (exclusively in the inferior occipital gyri for the 10 Hz frequency). For some participants, these oscillations extended further into the parietal and temporal cortices (see example maps on Figure 4c and average magnitudes across the brain on Figure 4d). Significant oscillations were also found in the later time window (150–210 ms) at frequencies from 6.8 Hz to 13.5 Hz for a minimum of four participants (population prevalence MAP = 0.79, 96% HPDI = [0.36, 0.98]). These were generally observed in slightly higher-level areas, in the middle, inferior, and anterior occipital gyri (Figure 4c,e). Interestingly, significant oscillations were observed in a majority of participants even at the relatively high 30.4 Hz frequency (3/5 participants; population prevalence MAP = 0.58, 96% HPDI = [0.18, 0.90]). These results are, to our knowledge, the first direct demonstration of oscillatory *sampling* in the brain. Such sampling oscillations between 7 and 17 Hz are consistent with findings from previous studies of perceptual oscillations (Blais et al., 2013; Fiebelkorn et al., 2013; VanRullen et al., 2007; see VanRullen, 2016, for a review).

Importantly, this oscillatory sampling must not be confused with oscillatory processing: while the latter refers to information that is processed non-continuously, in an oscillating manner, the former refers to information received at successive moments being processed and not processed, in a periodic, alternating way (Figure 4a). Oscillations in processing have also been documented in the brain (e.g., Rousselet et al., 2007; Schyns et al., 2011; Smith et al., 2006). However, which type of oscillations — either sampling or processing — is more prevalent remains unknown. To quantify this, we fitted models that explained all the activity within each map with either processing oscillations or sampling oscillations (while accounting for the larger processing delays for later stimulus moments) and assessed which model explained the activity the best (amplitudes and phases were free parameters; see Methods). Sampling oscillations explained features except the right eye in the expression task (3/5 participants; population prevalence MAP = 0.59, 96% HPDI = [0.20, 0.90]; within-subject randomization test; *p* < .05, two-tailed, FWER-corrected). This was observed in brain regions similar to the ones with significant oscillations uncovered in the previous analysis: the occipital poles, and inferior, middle and superior occipital gyri (Figure 4f,g). For the mouth in the expression task, all participants exhibited the effect in the right superior occipital gyrus (population prevalence MAP = 1.00, 96% HPDI = [0.57, 1.00]). The reverse result (processing oscillations explaining activity better than sampling oscillations) was a much rarer occurrence, never present in more than one participant for any brain source, task and face feature (Extended Data Figure 4-1). These results suggest that successive cycles of endogenous oscillations in early visual areas are allocated sequentially to information received at successive moments, instead of all coding information received at the same moment(s).

## Discussion

We developed a novel method to isolate the processing of specific visual features received at specific moments across the brain. This allowed us to reveal a new aspect of visual processing and characterize more fully the dynamics of object recognition. We first showed that processing is highly dependent on when information is received on the retina. Many different processes could be responsible for such variance in activity across stimulus moments. These include bottom-up mechanisms such as intrinsic neural oscillations, temporal integration, priming and adaptation, but also top-down predictive and attentional mechanisms. These different processes are likely to result in distinct patterns of activity across stimulus moments (Caplette et al., 2020).

We identified three main types of patterns in our study. In the first type, mostly observed in the earliest visual areas, information from most stimulus moments was processed around 100 ms after it was received. This is consistent with an early and automatic bottom-up processing of all incoming visual information by sensory brain areas. Moreover, this processing oscillated as a function of stimulus moment. This suggests that the processing was modulated by endogenous neural oscillations in these brain regions. In patterns of the second type, predominant in both similar and higher-level occipital areas, only early stimulus moments were processed, at a latency of about 170 ms. These early stimulus moments may be favored because the total amount of useful information that will appear in a given trial is uncertain (Nienborg & Cumming, 2009). Why this occurs around 170 ms is unclear but could be related to the N170 component, which is typically observed around that latency in similar brain areas. There is some evidence that a gating of some information prior to further processing happens around that component’s latency (Caplette et al., 2020; Zhan et al., 2019). Finally, we identified a third type of pattern: in these occurrences, information from most stimulus moments was processed around 440 ms and in higher-level regions such as the inferior occipital gyrus and the parietal cortex. These maps likely represent higher-level processing occurring in later brain areas of the visual hierarchy, prior to the preparation of the motor action. This processing might be related to evidence accumulation and the computation of the decision (Hanks et al., 2015; Woolnough et al., 2020).

Importantly, our method allowed us to reveal and quantify *sampling* oscillations in the brain (i.e., different stimulus moments being processed to a different degree, in a periodic fashion) for the first time. We showed that such oscillations occur mostly in early visual areas, at most analyzed frequencies, in the theta, alpha and beta bands. Perceptual oscillations in the theta and alpha range had previously been observed during sustained attention tasks. In some of these studies, accuracy during visual detection and discrimination tasks seemed to fluctuate at frequencies between 4 and 10 Hz (e.g., Fiebelkorn et al., 2013; Landau & Fries, 2012; Senoussi et al., 2019). In others, the prestimulus phase of alpha band EEG activity modulated either perceptual behavior (e.g., Busch & VanRullen, 2010; Helfrich et al., 2018; Fiebelkorn et al., 2018) or neural correlates of perception (e.g., Busch & VanRullen, 2010; Hanslmayr et al., 2013). We also observed significant low beta oscillations between 13 and 17 Hz: these frequencies are similar to those reported by Blais et al. (2013) who behaviorally assessed the frequencies at which information was sampled by human subjects in a face recognition task. Other studies also reported behavioral oscillations around 13 Hz in motion perception and face recognition tasks (Macdonald et al., 2014; VanRullen et al., 2005; Vinette et al., 2004). In general, the presence of sampling oscillations in these frequency bands is consistent with findings from many previous behavioral and electrophysiological studies about perceptual oscillations (VanRullen, 2016; although several other studies failed to uncover similar findings, see Keitel et al., 2022; Ruzzoli et al., 2019; van der Werf et al., 2021).

Interestingly, sampling oscillations in the brain were more prevalent than processing oscillations (i.e., oscillations in the processing of information received at a given moment, e.g., Schyns et al., 2011; Smith et al., 2006). This suggests that successive cycles of endogenous oscillations in early visual areas are allocated to stimulus information incoming at successive moments, rather than responding to information from the same stimulus moment. In other words, instead of being reflected as an oscillatory processing of some snapshot of the visual field, brain oscillations seem to lead to a rhythmic sampling of the continuous flow of incoming visual information. This coding scheme makes sense since it allows the visual cortex to be continuously updated with the most recent sensory information. This should be especially helpful to the organism if sensory stimuli were to change during the fixation.

Note that our statistical procedure is not affected by the same problems as most studies of rhythmic sampling (as recently argued by Brookshire, 2022; see also Kienitz et al., 2021). Indeed, we are not using a time-shuffling method (but rather a trial-shuffling one) and our empirical null distribution has the same aperiodic temporal structure as our observed data. Therefore, our statistical threshold should be appropriate for all frequencies. That being said, significant Fourier spectrum peaks do not necessarily indicate the presence of true *oscillations*, this term usually referring to temporally extended periodic activity. In particular, whether the activity at the 6.8 Hz frequency – the lowest analyzed frequency – can be considered truly oscillatory is uncertain, and we therefore recommend caution when interpreting this result. Furthermore, we must stress that our findings of rhythmic sampling in the brain do not inform us about *perceptual* (behavioral) rhythmic sampling, that is, the idea that human perception is fundamentally discrete or cyclic (see Blais et al., 2013; Landau & Fries, 2012; Keitel et al., 2022; VanRullen, 2016). Future studies should examine the link between behavioral sampling and the rhythmic sampling in the brain that we have uncovered.

In summary, we developed a method to disentangle information sampling and processing in the brain in a fine-grained manner. We showed that the moment at which information is received greatly modulates information processing, that these modulations are varied across the brain, and that sampling is oscillatory in early visual areas. Specifically, we showed that these oscillations in sampling occur early on during visual processing, at multiple frequencies peaking in the theta and low beta bands, and that they are more prevalent than the previously investigated oscillations in processing. These findings further our understanding of the role of brain oscillations in visual processing. More generally, our results also show that considering the temporal extent of stimuli, in addition to the temporal dimension of brain activity, is necessary to understand visual recognition more fully. Other phenomena could be investigated in future experiments using this paradigm, including temporal integration, attentional sampling and visual routines.

## Acknowledgements

We wish to thank Nicolas Dupuis-Roy and Robin Ince for help with the experimental design and analysis. We also thank the five participants without whom this project would not have been possible. Finally, we thank our funding sources: NSERC (Discovery Grant and Alexander-Graham-Bell Doctoral Scholarship), FRQNT (Postdoctoral Fellowship) and University of Montreal (Doctoral Scholarship).

## Multimedia Legends

**Video 1.** Example of a random stimulus.

**Video 2.** Same as Video 1, slowed down 10 times.

## Extended Data Figure Legend

**Extended Data Figure 4-1.** Sampling oscillations vs processing oscillations model fit, for each face feature, for the gender (a) and expression (b) tasks. The color scale refers to the number of participants for which the processing oscillations model fit better (within-subject randomization test, *p* < .05, two-tailed, FWER-corrected) and the Bayesian MAP of the estimated population prevalence. The maximum number of participants across the cortex is indicated below for each feature and task. A better fit of the processing oscillations model never occurred for more than one participant (see Figure 4 for the reverse effect).

## Author Contributions (CRediT)

L.C.: Conceptualization, methodology, software, project administration, investigation, data curation, formal analysis, visualization, writing – original draft, writing – review & editing.

K.J.: Methodology, supervision, validation, writing – review & editing.

F.G.: Conceptualization, methodology, supervision, validation, funding acquisition, writing – review & editing.

